# Declining Genetic Polymorphisms of the C-terminus Merozoite Surface Protein-1 Amidst Increased *Plasmodium knowlesi* Transmission in Thailand

**DOI:** 10.1101/2024.06.27.600943

**Authors:** Parsakorn Tapaopong, Sittinont Chainarin, Abdulrohman Mala, Arnuwat Rannarong, Nikom Kangkasikorn, Teera Kusolsuk, Wanlapa Roobsoong, Liwang Cui, Wang Nguitragool, Jetsumon Sattabongkot, Sirasate Bantuchai

**Affiliations:** Mahidol Vivax Research Unit, Faculty of Tropical Medicine, Mahidol University, Bangkok, Thailand; Department of Molecular Tropical Medicine and Genetics, Faculty of Tropical Medicine, Mahidol University, Bangkok, Thailand; Ruso Hospital, Ruso, Narathiwat, Thailand; Vector-Borne Disease Control Center 3.2, Nakhon Sawan, Thailand; Department of Helminthology, Faculty of Tropical Medicine, Mahidol University, Bangkok, Thailand; Division of Infectious Diseases and International Medicine, Department of Internal Medicine, Morsani College of Medicine, University of South Florida, Tampa, Florida, USA

**Author notes:** Corresponding authors: Correspondence to Mahidol Vivax Research Unit, Faculty of Tropical Medicine, Mahidol University, Bangkok, Thailand. Email address (S. Bantuchai).

## Abstract

Recent reports from Thailand reveal a substantial surge in *Plasmodium knowlesi* cases over the past decade, with a more than eightfold increase in incidence by 2023 compared to 2018. This study investigates temporal changes in genetic polymorphism associated with the escalating transmission of *P. knowlesi* malaria in Thailand over the past two decades. Twenty-five *P. knowlesi* samples collected in 2018–2023 were sequenced for the 42-kDa region of *pkmsp1* and compared with 24 samples collected in 2000–2009, focusing on nucleotide diversity, natural selection, recombination rate, and population differentiation. Seven unique haplotypes were identified in recent samples, compared to 15 in earlier samples. Nucleotide and haplotype diversities were lower in recent samples (π = 0.016, Hd = 0.817) than in earlier samples (π = 0.018, Hd = 0.942). Significantly higher synonymous substitution rates were observed in both sample sets (d_S_ – d_N_ = 2.77 and 2.43, p < 0.05), indicating purifying selection and reduced genetic diversity over time. Additionally, 8 out of 17 mutation points were located on B-cell epitopes, suggesting an adaptive response by the parasites to evade immune recognition. Population differentiation analysis using the fixation index (F_st_) revealed high genetic differentiation between parasite populations in central and southern Thailand or Malaysia. Conversely, the relatively lower F_st_ value between southern Thailand and Malaysia suggests a closer genetic relationship, possibly reflecting historical gene flow. In conclusion, our findings highlight a decline in genetic diversity and evidence of purifying selection associated with the recently increased incidence of *P. knowlesi* malaria in Thailand. The minor genetic differentiation between *P. knowlesi* populations from southern Thailand and Malaysia suggests a shared recent ancestry of these parasites and underscores the need for coordinated efforts between the two countries for the elimination of *P. knowlesi*.

## INTRODUCTION

The emergence of *Plasmodium knowlesi* as the fifth human malaria parasite in Southeast Asia (1, 2) has grasped substantial attention within the malaria research community. The incidence of *P. knowlesi* malaria exhibits enormous geographical variations, with the highest rates recorded in Malaysia (1). Despite its lower prevalence compared to *P. falciparum* and *P. vivax* (3, 4, 5, 6), *P. knowlesi* poses a considerable public health threat in Southeast Asia (1, 7, 8). Recently, Thailand observed a significant increase in *P. knowlesi* cases (Fig 1A) (9, 10). However, whether this surge was due to parasite introduction or increased transmission of existing parasite reservoirs remains uncertain. Notably, numerous cases of co-infection involving *P. knowlesi* and other human malaria species have been documented, especially in the southern provinces of Thailand (3, 4, 5). Given *P. knowlesi*’s 24-hour intraerythrocytic cycle, infections can lead to rapid increases in parasitemia and severe illness, potentially resulting in fatalities if treatment is delayed (1, 8, 11). *P. knowlesi* primarily infects long-tailed macaques (*Macaca fascicularis*) and pig-tailed macaques (*Macaca nemestrina*), which widely habitate Southeast Asia (12). While transmission to humans typically occurs through mosquito bites originating from infected macaques, experimental evidence suggests the potential for human-to-human transmission (13).

**Fig 1.**
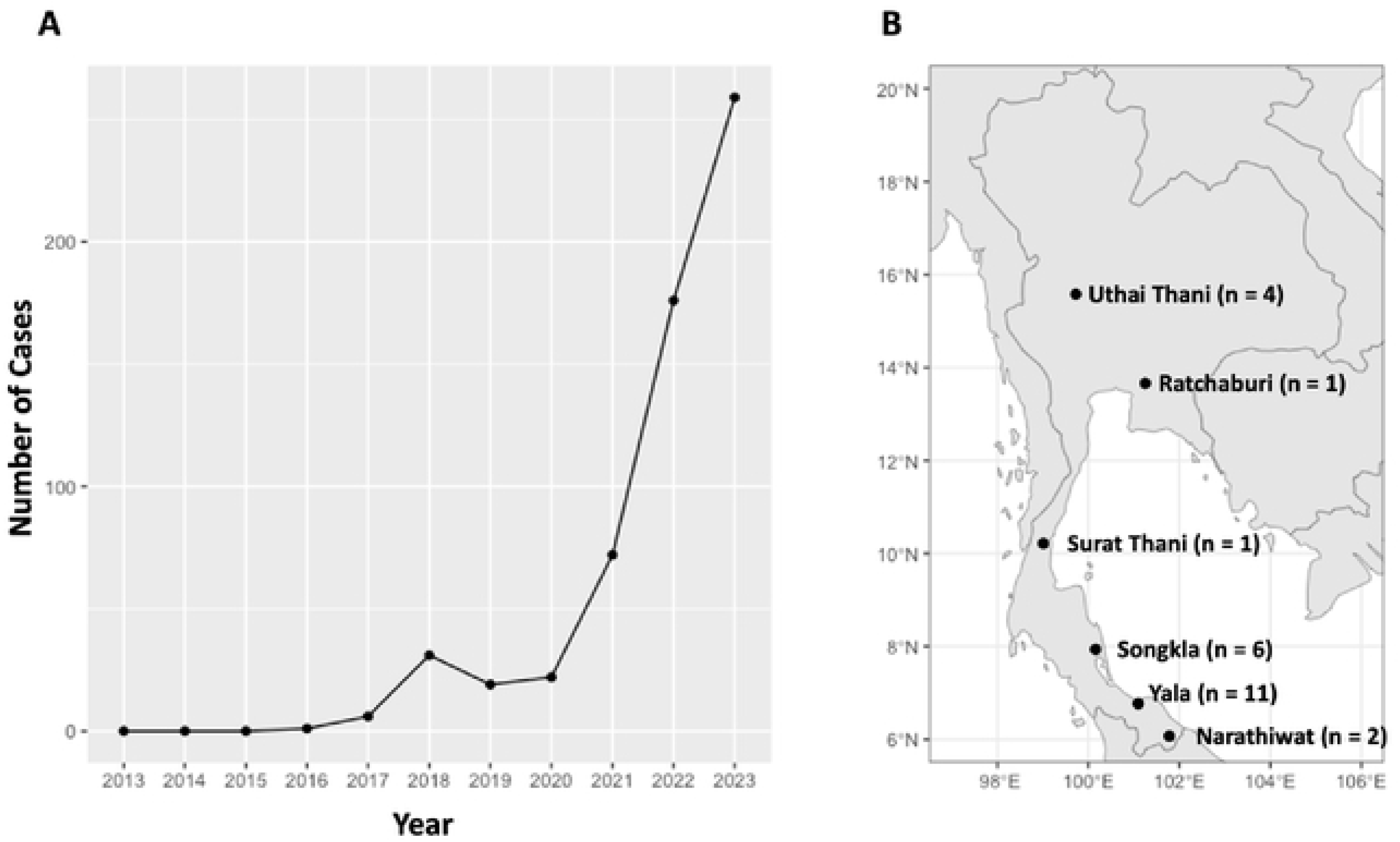
Increase in *P. knowlesi* incidents over the past decade and sample collection sites in Thailand. (A) The line chart illustrates the escalating number of malaria incidents attributed to *P. knowlesi* in Thailand frorn 2013 to 2023. (B) The map highlights the collection sites of parasite isolates, with “n” denoting the number of cases collected fro1n each province.

High levels of genetic variation have been identified within orthologous antigens of known vaccine candidates in *P. knowlesi*, including Duffy binding protein (*pkdbp*), normocyte binding proteins xa and xb (*pknbpxa* and *pknbpxb*), and merozoite surface protein-1 (*pkmsp1*). This extensive variation presents a significant challenge to malaria vaccine development (14, 15, 16, 17). Consequently, assessing the degree of genetic polymorphisms, natural selection, and population structure of these antigenic markers is crucial not only for vaccine development but also for gaining insights into transmission dynamics and population genetics of *P. knowlesi*. However, limited information is available regarding genetic variation within this parasite species, primarily due to its low prevalence and restricted access to sample collection. Addressing these knowledge gaps is essential for advancing our understanding of *P. knowlesi* biology and informing the development of effective control strategies.

Merozoite surface protein-1 (*msp1*) has been a key marker for studying genetic diversity across various *Plasmodium* species (4, 5, 14, 18, 19, 20). In *Plasmodium*, *msp1* undergoes complex processing, initially synthesized as a 200 kDa precursor and later cleaved into four fragments during merozoite maturation (21). Notably, the C-terminal 42-kDa fragment is further cleaved into *msp1_33_* and *msp1_19_* during merozoite invasion, with *msp1_19_* remaining on the merozoite surface (22). Due to its significant immunogenicity, the 42-kDa fragment is a potential vaccine candidate (23). *In vitro* studies have shown that antibodies against *msp1_42_* and *msp1_19_* can inhibit *P. falciparum* merozoite invasion (24, 25). However, extensive genetic polymorphisms in *msp1_42_*among global isolates of *P. falciparum* and *P. vivax* pose challenges for vaccine development (26, 27, 28). Although the sequences of *pfmsp1* and *pvmsp1* have been extensively characterized (29, 30), the sequence of *pkmsp1* remains relatively understudied. Recent investigations have revealed high genetic variation in *pkmsp1_42_*in Malaysia (14, 19), analyzing intra- and inter-population diversities of *pkmsp1* from different geographical isolates, including Malaysian Borneo, Peninsular Malaysia, and Thailand (20). Despite this, limited studies have been performed on *pkmsp1* in Thailand, with only 24 isolates (12 from humans and 12 from monkeys, collected from 2000 to 2009) analyzed to date (4, 5). Given the importance of *pkmsp1* as an immunogenic antigen, there is a notable gap in understanding its genetic diversity within Thai clinical isolates.

To address this gap, we collected additional *P. knowlesi* samples during 2018 –2023, coinciding with a significant increase in *P. knowlesi* malaria incidence (4, 5). Our primary objective was to investigate the temporal changes in genetic polymorphisms associated with the increased transmission intensity of *P. knowlesi* malaria observed in Thailand over the past decade. Additionally, we aimed to assess inter-population diversities of *pkmsp1* among isolates from Thailand, Malaysian Borneo, and Peninsular Malaysia. This study sheds light on the evolving genetic landscape of *P. knowlesi* in this region, contributing to a better understanding of its transmission dynamics and potential implications for malaria control strategies.

## METHODS

### Ethics statement

The use of parasite DNA samples and associated data in this study received ethical approval from the Ethics Committee of the Faculty of Tropical Medicine, Mahidol University, Bangkok (Approval numbers: MUTM2018-016-05 and MUTM2017-078-05).

### Samples collection and confirmation of *P. knowlesi* infection in human blood samples

Parasite DNA was extracted from whole blood or dried blood spots (DBS) collected from malaria-positive patients aged 15 to 80 in various provinces of Thailand, including Songkla (n = 6), Yala (n = 11), Narathiwat (n = 2), Surat Thani (n = 1), Ratchaburi (n = 1), and Uthai Thani (n = 4), during the period from 2018 to 2023 (Fig 1B). Detection of *P. knowlesi* infections was conducted using nested PCR assays targeting 18S rRNA genes, following a previously published protocol (10). All DNA samples were stored at −20°C and archived at the Mahidol Vivax Research Unit (MVRU) until use.

### Gene amplification, purification, and sequencing of *pkmsp1_42_*

Due to the limited parasite DNA from dried blood spots in some samples, the REPLI-g Mini Kit (Qiagen, Germany) was employed for whole genome amplification (WGA) of *P. knowlesi*-infected samples, following the manufacturer’s protocol. The REPLI-g amplified DNA was stored at −20°C until further use. PCR amplification of the *pkmsp1_42_* fragment was conducted in a 25 µl reaction volume, comprising 3 µl of DNA template, GoTaq Green master mix (Omega, USA), and 0.4 µM forward and reverse primers (MSP-1_3934_F: 5’-CGCTACAGGCCTATGAGGAA-3’ and MSP- 1_5431_R: 5’-AGGAAGCTGGAGGAGCTACA-3’). The PCR conditions consisted of an initial cycle at 94°C for 5 min, followed by 35 cycles of 94°C for 45 sec, 61°C for 45 sec, and 72°C for 1 min, with a final extension step at 72°C for 10 min. The amplified products were visualized on 1.5% agarose gels and subjected to standard dye terminator sequencing (Macrogen, Korea) using MSP-1_3934_F as the sequencing primer.

### Retrieval of additional sequences

For comparative analysis, homologous *pkmsp1_42_* sequences of *P. knowlesi* collected between 2000 and 2009 by Putaporntip et al., 2013 (5) were retrieved from the NCBI GenBank database (n = 24, accession numbers JF837339 – JF837353, JX046791 – JX046798, and DQ220743) to assess genetic diversity between recent and former populations. Additionally, *pkmsp1_42_* sequences of isolates from Malaysia, as collected by Yap et al., 2017 (14) and Noordin et al., 2023 (31), were obtained from the NCBI GenBank database (n = 36; accession numbers KX881363 – KX881371, KX894505 – KX894507, and ON926539 – ON926561) for geographical comparison.

### DNA sequence polymorphism analysis

The 25 *pkmsp1_42_* sequences were aligned with the reference sequence of the *P. knowlesi* H strain (PlasmoDB accession number PKNH_0728900) using SnapGene Viewer version 6.2.1, with manual trimming of low-quality sequences. DnaSP v5.1 software was employed to calculate various parameters, including the number of segregating sites (S), the number of mutations (η), the average number of nucleotide differences (k), the number of haplotypes (H), haplotype diversity (Hd), and nucleotide diversity (π), along with their corresponding standard deviations (SD). A Z-test of selection (p < 0.05) was computed for the difference between synonymous and nonsynonymous substitutions (d_S_ - d_N_) based on the alternative hypothesis (H_A_: d_S_ > d_N_) using the Nei and Gojobori method (32) with Juke and Cantor correction, and bootstrap variance estimation of 1000 replicates in MEGA v10.2.6 software (33). Additionally, Tajima’s D and Fu and Li’s D* and F* tests were performed to assess deviation from neutrality. The minimum number of recombination events (Rm) and linkage disequilibrium (LD) were computed in terms of R^2^ and D′ indices related to nucleotide distance using DnaSP v5.1 software. Twenty-four additional *pkmsp1_42_*sequences from Thailand in 2000 – 2009 were retrieved from GenBank and combined with the 25 sequences from this study for temporal change in genetic diversity analyses. A haplotype network for *pkmsp1_42_* was generated using the median-joining method in the POPART v10.1 software (34). To evaluate the degree of genetic differentiation based on *pkmsp1_42_*, Wright’s fixation (F_st_) statistics indexes (35) were calculated with DNA sequences between central and southern Thailand or Malaysia isolates using DnaSP v5.1 software. Linear B-cell epitopes were predicted from the 42 kDa region of *pkmsp1* using Bepipred under the default threshold score of 0.5 (http://tools.iedb.org/main/bcell/) (36). Furthermore, the positions of nonsynonymous changes on predicted B-cell epitopes were visualized on the crystal structure of *pkmsp1_42_* (UniProt Accession Number: K7QLU5) using PyMOL software v2.5.2.

## RESULTS

### Nucleotide polymorphism and amino acid changes

From 49 samples collected in Thailand (25 from 2018 – 2023 and 24 from 2000 – 2009), the 456 bp fragment (nucleotide position 4450 to 4905) of the *pkmsp1_42_* was sequenced. Sequence alignment against the reference strain H revealed that the 25 recent samples contained 19 single nucleotide polymorphisms (SNPs) in the *pkmsp1_42_*, with 10 SNPs being synonymous and 9 nonsynonymous. Among these SNPs, 3 were singletons and 16 were parsimony informative. Conversely, the 24 earlier samples had 29 SNPs in the *pkmsp1_42_*, with 13 synonymous and 16 nonsynonymous SNPs. Of these SNPs, 8 were singletons and 21 were parsimony informative (supplementary material S1 and S2).

Among the 17 amino acid changes identified in all samples, 8 were shared between the earlier and recent samples. Eight mutations (D1495N, I1547L, I1556T, K1584N, Q1594H, Q1596P, E1603Q, K1629N) were exclusively present in the earlier samples, and one mutation (A1588S) was exclusively found in the recent samples. Upon stratification by the sampling period, differential mutation frequencies were observed for N1513S (16.7% and 4%), N1566K (25% and 24%), Q1574K (41.7% and 8%), G1592S (20.8% and 32%), E1593G (16.7% and 32%), A1599T (8.3% and 32%), Q1600E (25% and 32%), and M1624I (8.3% and 32%) in the earlier and recent periods, respectively (Fig 2, supplementary material S4).

**Fig 2.**
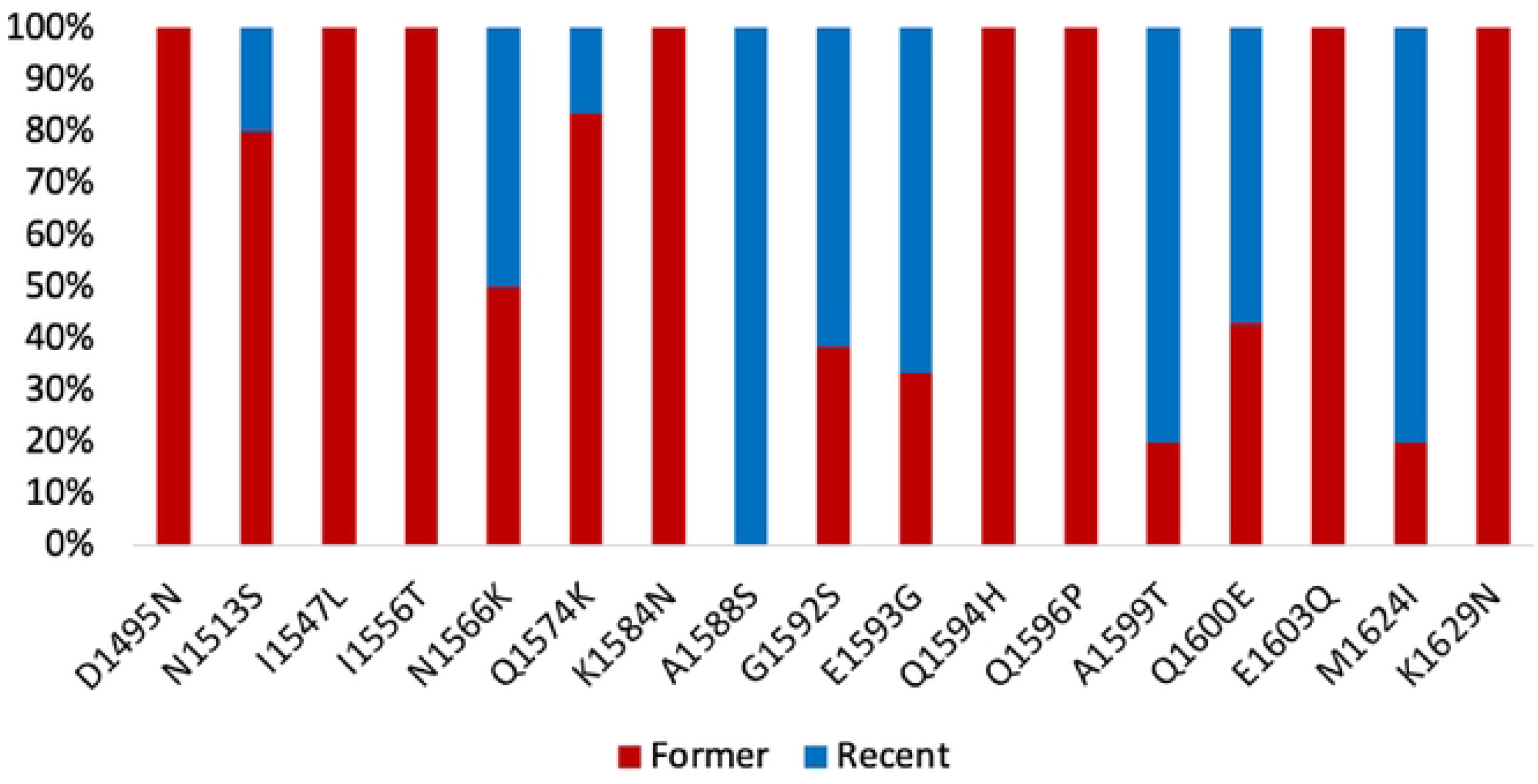
Frequencies of amino acid changes detected in *pkmsp1 _2_* among Thailand isolates. The red and blue segments of the columns depict the propo1tion of amino acid changes observed in the former period (2000 - 2009) and the recent period (2018 - 2023), respectively.

The nucleotide sequence of the 42-kDa region of *pkmsp1* was translated into an amino acid sequence and analyzed using Bepipred to predict linear B cell epitopes. Seven residues or peptides within the internal stretch of the *pkmsp1_42_* sequence (NITDMLDSRLKKRNY, DSELNPFKYSSSGEYIIKDPYKLLDLEQKKKLLGS, EY, NKMGELYKQ, NKMGELYKQ, KEYESLVN, and E) were identified as potential B-cell epitopes (Fig 3A). Analysis revealed that 8 of the 17 mutations (N1513S, N1566K, Q1574K, G1592S, E1593G, A1599T, Q1600E, and M1624I) were located on B-cell epitopes and were shared between the earlier and recent samples (Fig 3B), suggesting an adaptive response by the parasites to evade immune recognition.

**Fig 3.**
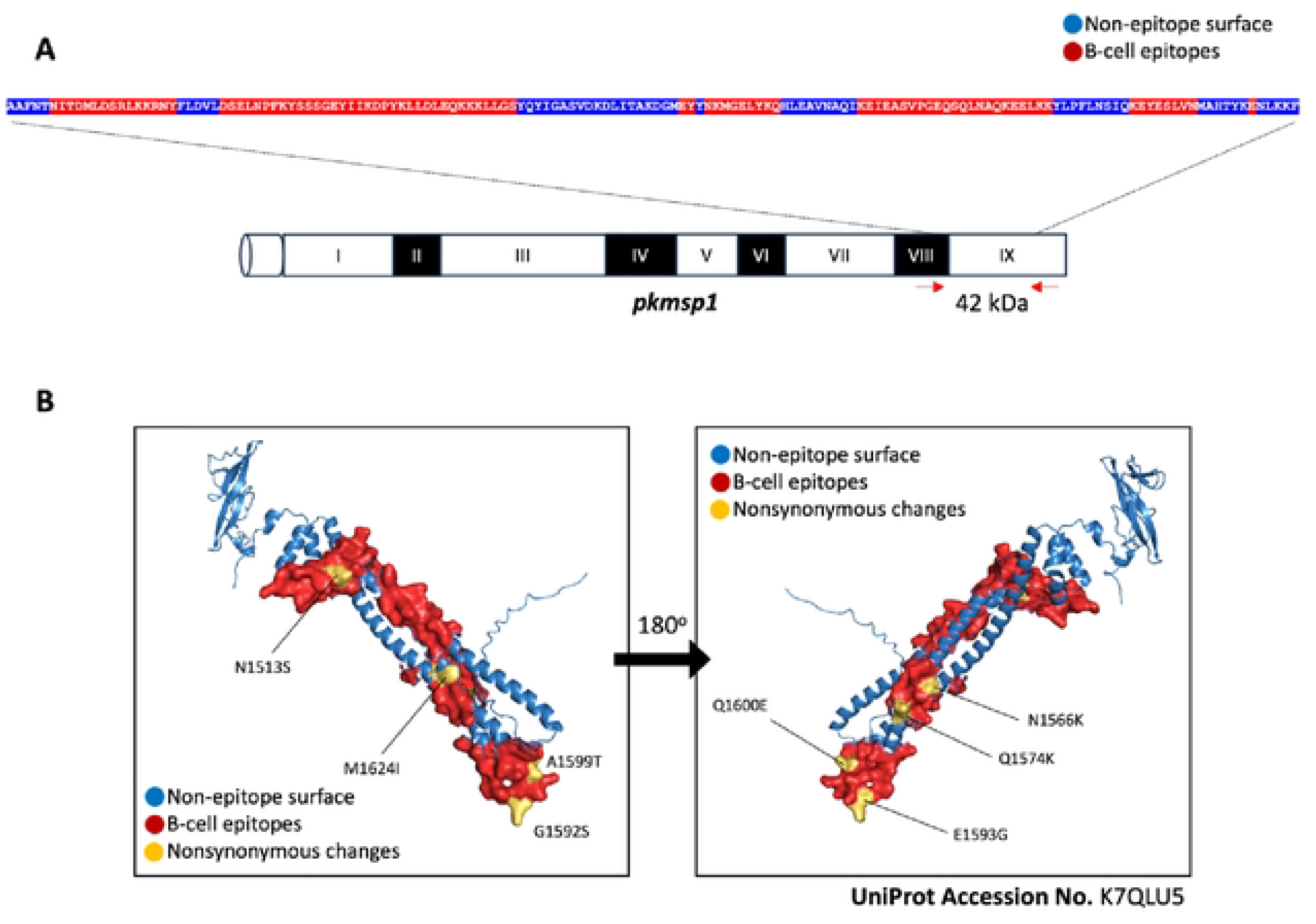
Prediction of B-cell epitopes on *pkmsp1_42_* and location of nonsynonymous changes. (A) The peptide sequence of the 42 kDa region of *pkmspl* is shown, with predicted potential B-cell epitopes highlighted in red residues and non-epitope surface residues in blue within the internal stretch of the sequence. (B) Based on the structure of the 42 kDa region of *pkmspl* (UniProt Accession No. K7QLUS), the locations of nonsynonymous changes of each amino acid shared between former and recent isolates found in *pkmspl* are labeled in yellow. Additionally, the locations of predicted B-cell epitopes and non-epitope surfaces are colored in red and blue, respectively.

### Temporal changes in genetic diversity, haplotypic relationship, and natural selection

We analyzed the genetic diversity of the 42-kDa region of *pkmsp1* and identified 7 and 15 unique haplotypes of *pkmsp1_42_* in the recent and earlier samples, respectively. Notably, we observed lower nucleotide diversity (π) and haplotype diversity (Hd) in the recent samples (π = 0.016, Hd = 0.817) than in the earlier samples (π = 0.018, Hd = 0.942). Additionally, the average number of nucleotide differences (k) was lower in the recent samples, suggesting a decline in genetic heterogeneity over the past decade amidst higher transmission intensity of *P. knowlesi* in Thailand (Table 1).

**Table 1.**
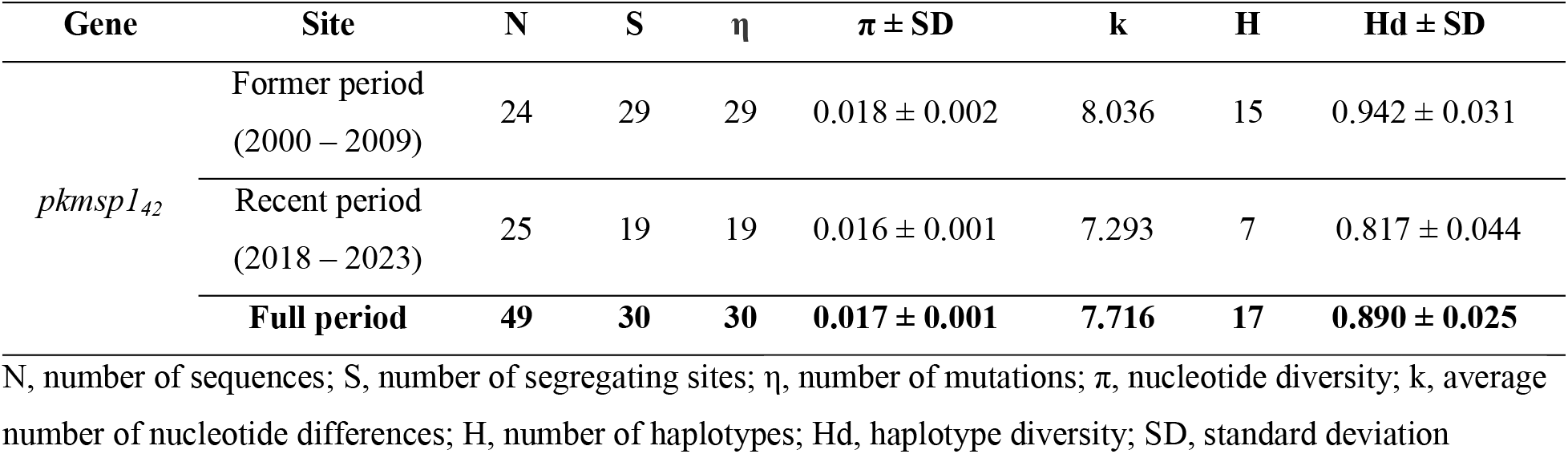
Genetic diversity of *pkmsp1_42_*.

| Gene | Site | N | S | $\eta$ | $\pi \pm \text{SD}$ | k | H | Hd ± SD |
| --- | --- | --- | --- | --- | --- | --- | --- | --- |
| *pkmsp1*<sub>42</sub> | Former period<br>(2000 – 2009) | 24 | 29 | 29 | 0.018 ± 0.002 | 8.036 | 15 | 0.942 ± 0.031 |
| | Recent period<br>(2018 – 2023) | 25 | 19 | 19 | 0.016 ± 0.001 | 7.293 | 7 | 0.817 ± 0.044 |
|  | **Full period** | **49** | **30** | **30** | **0.017 ± 0.001** | **7.716** | **17** | **0.890 ± 0.025** |

N, number of sequences; S, number of segregating sites; $\eta$ , number of mutations; $\pi$ , nucleotide diversity; k, average number of nucleotide differences; H, number of haplotypes; Hd, haplotype diversity; SD, standard deviation

A haplotype network was constructed using the 49 sequences (Fig 4). The sequences clustered into 17 haplotypes, with each haplotype separated by one to six mutational steps. Notably, ten haplotypes were represented by a single sequence, most of which (H3, H4, H5, H6, H8, H11, H12, H13, H14, and H16) were present in the earlier samples. Haplotype 9 was the most predominant, consisting of 11 isolates. Five haplotypes (H1, H2, H7, H9, and H15) were shared between the recent and earlier samples, whereas only two were specific to the recent samples. This suggests that most haplotypes in the recent population were inherited from the earlier population, with rare new haplotypes emerging. Additionally, four haplotypes (H3, H6, H7, and H12) were shared between Thailand and Malaysia, indicating some genetic similarity between isolates from these two regions (Fig 5).

**Fig 4.**
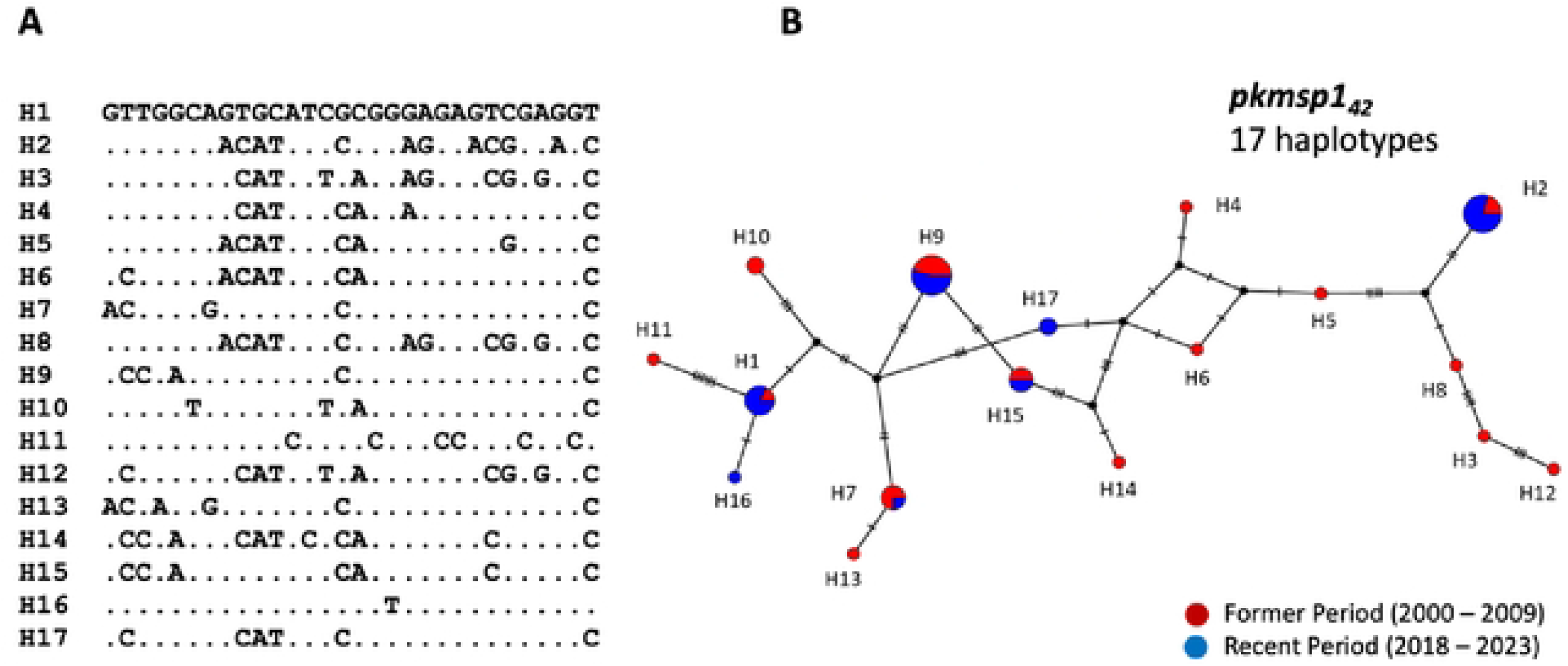
Haplotypic relationship of parasite population between forn1er and recent period. (A) Identification of seventeen unique haplotypes of *pkmspl_2_* fro1n a total of 49 *P. knowlesi* isolates in Thailand. (B) Construction of a haplotype network using the Median-joining algorith1n, illustrating the relationship between the 17 haplotypes (H1-H17) of *pkmspl _42_* fro1n the former population (in red) and recent population (in blue). Each circle represents a haplotype, with the size of the circle indicating the haplotype frequency. The distance between circles represents the degree of genetic relatedness, with the number of nucleotide-base changes labeled as hatch marks.

**Fig 5.**
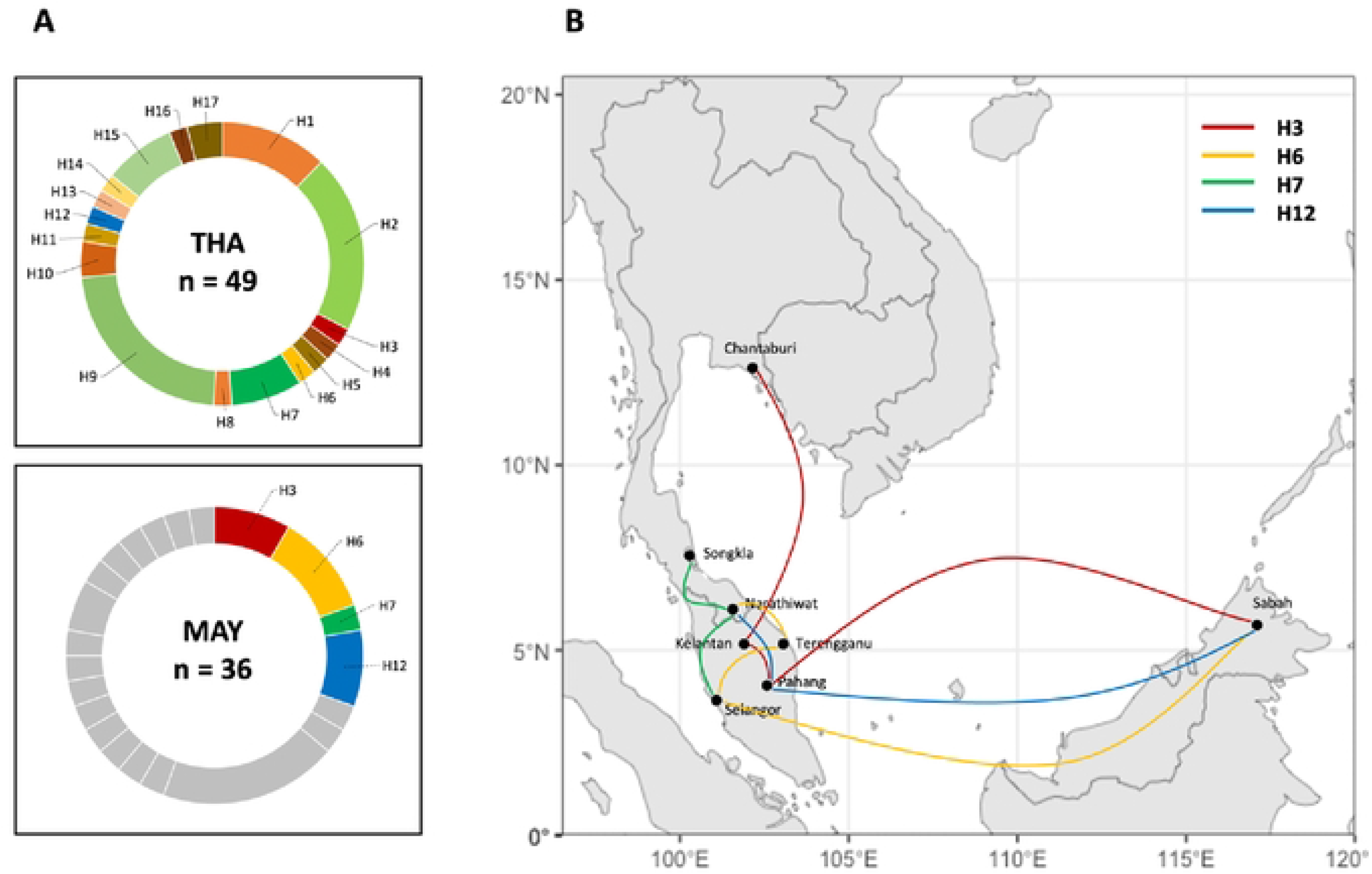
Regional distribution of shared haplotypes between Thailand and Malaysia. (A) Donut charts depict the frequencies of shared haplotypes labeled with geographic origins: THA for Thailand and MAY for Malaysia, where “n” represents the nu1nber of analyzed isolates. (B) The regional distribution of the four haplotypes shared between Thailand and Malaysia is illustrated among the collection sites.

Non-statistically significant values were obtained from several neutrality tests (Tajima’s D, Fu and Li’s D*, and F*). Additional investigations were conducted to elucidate the evolutionary processes shaping *pkmsp1_42_* genetic variations. Specifically, the Z-test for the difference between rates of synonymous and nonsynonymous substitutions (d_S_–d_N_) was performed to assess natural selection. Significant Z-test results were observed in both earlier (d_S_–d_N_ = 2.77) and recent periods (d_S_–d_N_ = 2.64), indicating an excess of synonymous changes over nonsynonymous changes in *pkmsp1_42_* (Table 2). This observation suggests that purifying selection is likely driving the evolution of *P. knowlesi* in Thailand, favoring the maintenance of functional integrity within the 42-kDa region of the *pkmsp1* gene.

**Table 2.** Neutrality tests and recombination of *pkmsp1_42kDa_*.

| Gene | Site | $d_S - d_N$ | D | D* | F* | Rm |
| --- | --- | --- | --- | --- | --- | --- |
| *pkmspI*<sub>42</sub> | Former period<br>(2000 – 2009) | 2.77* | 0.131 | 0.022 | 0.065 | 6 |
|  | Recent period<br>(2018 – 2023) | 2.43* | 1.616 | 0.682 | 1.134 | 2 |
|  | **Full period** | **2.64\*** | **0.490** | **-0.567** | **-0.230** | **6** |

$d_S - d_N$ , Z-test for the difference between rates of synonymous substitution ( $d_S$ ) and nonsynonymous substitution ( $d_N$ ); D, Tajima's D test; D\*, Fu and Li's D\* value; F\*, Fu and Li's F\* value; Rm, minimum number of recombination events

### Recombination and linkage disequilibrium

Among the 49 samples analyzed, the recent samples showed a lower minimum number of recombination events (Rm=2) than the earlier samples (Rm=6). Additionally, analysis of the *pkmsp1_42_* fragments revealed significant linkage disequilibrium (LD) (p < 0.05, Fisher’s exact test), suggesting the presence of population structures among the parasites. This LD declined with increasing distance between pairs of polymorphic loci, suggesting the occurrence of recombination events within the *pkmsp1_42_* (Fig 6).

**Fig. 6:**
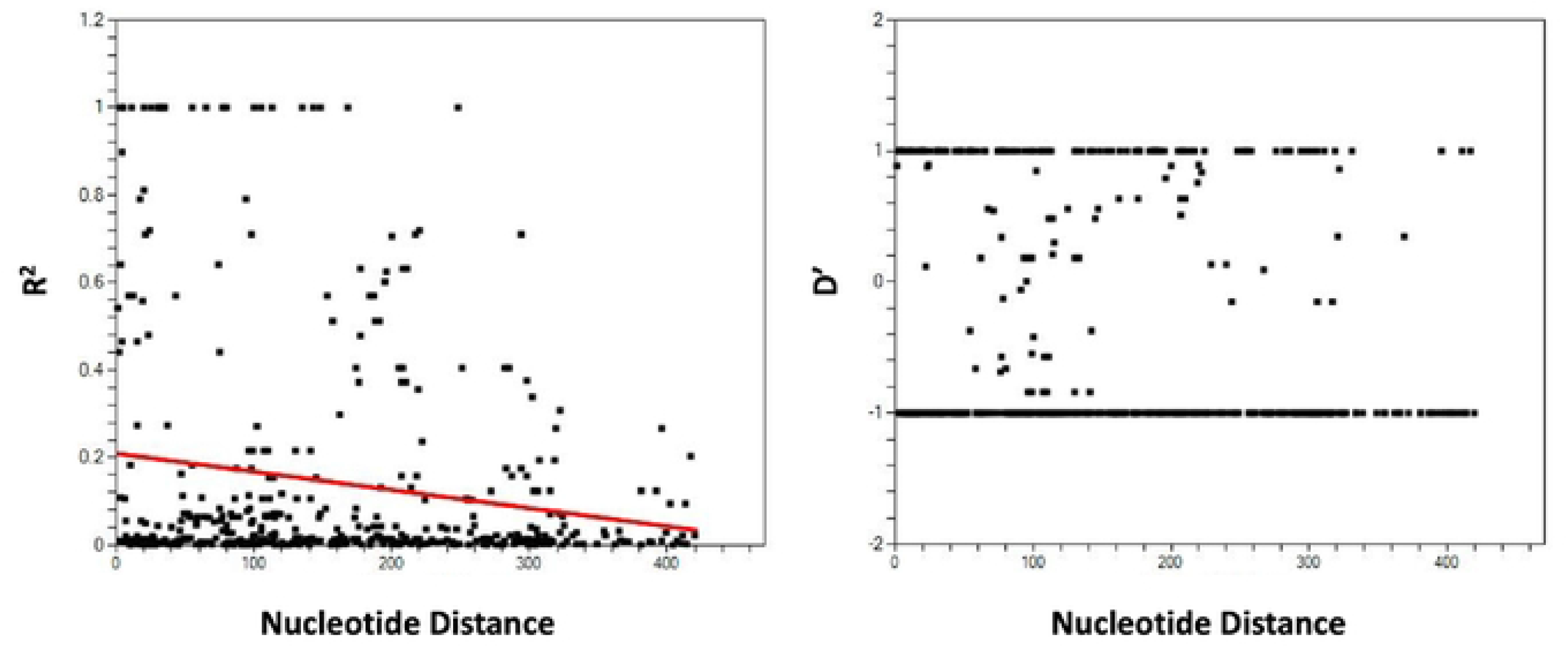
Linkage Disequilibrium (LD). The plot illustrates R^2^ and D’ against nucleotide distance using a two-tailed Fisher’s exact test for statistical significance. It represents the levels of meiotic recombination at different polymorphic sites between nucleotide variants within *pkmspl_42_*.

### Genetic differentiation in Thailand and Malaysia

To evaluate genetic differentiation across various endemic regions, we computed the fixation index (F_st_) using sequences from isolates in central Thailand (n = 11), southern Thailand (n = 38), and Malaysia (n = 36). Due to unequal sample sizes, we did not distinguish between Malaysian isolates from Peninsular and Borneo, as this could introduce sampling bias and affect F_st_ calculations. The F_st_ values exhibited considerable variation both within and between territories. Notably, comparisons between parasites from central and southern Thailand, as well as between central Thailand and Malaysia, revealed high levels of genetic differentiation for *pkmsp1_42_*, with F_st_ values of 0.3553 and 0.3231, respectively, suggesting lower levels of genetic exchange between central Thailand and the two southern populations. In contrast, we observed only moderate genetic differentiation between southern Thailand and Malaysia, with an F_st_ value of 0.1804 (supplementary S5 File). This finding is consistent with the geographic proximity of southern Thailand to Malaysia, implying higher levels of gene flow between these two regions.

## DISCUSSION

The increased incidence of *P. knowlesi* malaria in recent years has raised public health concerns in Thailand. The southern region of Thailand, which borders Malaysia, is considered a focal areas for *P. knowlesi* endemicity in Southeast Asia. Deforestation and urbanization may contribute to a shrinking habitat gap between humans and monkeys, facilitating shorter transmission distances for the disease via mosquitoes. Evidence of cross-transmission from monkeys to humans is supported by identical *pkmsp1* sequences found in both patients and macaques living in close proximity (37). While there is limited evidence of natural human-to-human transmission, experimental studies have shown that *P. knowlesi* can be transmitted from human to human via mosquitoes (13). The transmission of *P. knowlesi* in Southeast Asia is linked to the distribution of Leucosphyrus mosquitoes, known vectors of the parasite, which are predominantly found in the region. This limited distribution restricts the global spread of the disease (38).

Parasite genetic diversity is a potential indicator of transmission intensity (39). A decrease in transmission intensity has been associated with lower genetic diversity of *msp1*, as observed in northwestern Thailand and Mexico (40, 41). However, this trend is generally observed in low transmission settings, where factors such as random genetic drift, inbreeding, and reduced efficacy of natural selection contribute to the loss of genetic variation (42). Conversely, in high transmission areas, genetic diversity has remained high despite a decline in transmission intensity, as demonstrated by findings from Sri Lanka during 2003–2007 (43).

We observed a decline in genetic diversity within the 42 kDa region of *pkmsp1* over time, coinciding with increased transmission intensity. This decline is evidenced by lower levels of nucleotide and haplotype diversity among parasites collected in the recent period than in the earlier period. Notably, the earlier samples, which included 12 human and 12 monkey isolates, showed higher diversity in humans; further analysis focusing solely on human samples also revealed a similar decreasing trend in genetic diversity in the recent parasite population (supplementary material S6), consistent with the findings from the combined human and monkey samples.

The reduced diversity observed in the recent parasite population suggests a loss of variants over time, possibly due to the exclusion of alleles with deleterious mutations through negative purifying selection. This is supported by an excess of synonymous substitution rates (d_S_) over nonsynonymous substitution rates (d_N_) within the 42 kDa region of *pkmsp1*, indicating a preference for retaining H strain-like protein structures due to their essential functionality. The signatures of negative selection pressure and expansion of the population detected within conserved domains further suggest that the protein may be subject to strong functional constraints. Although statistical significance was not achieved, the negative values of neutrality tests (Fu and Li’s D* and F*) provide additional support for possible clonal expansion and indicate the presence of negative natural selection within the 42 kDa domain.

Approximately half of the 17 observed amino acid changes were not detected in the recent parasite population. This suggested that these mutations may confer reduced fitness to the parasites. Conversely, the remaining protein mutations, shared between the recent and former populations, were identified within the B-cell epitope, indicating host immune selection.

Upon red blood cell invasion, the 33 kDa segment is cleaved from the 42 kDa segment, subsequently binding to the protein inflammatory cytokine (S100P). This interaction attenuates the host’s inflammatory response and chemotactic activity (44). The 42 kDa region of *pkmsp1* has, therefore, garnered attention as a parasite protein, associated with chemotactic activity in host. Despite its potential, there has been limited genetic characterization of clinical isolates within this domain to assess population-level polymorphisms, which is crucial for evaluating vaccine feasibility. Although a degree of genetic diversity was observed in our study, the C-terminal domain IX (42 kDa) exhibited the lowest nucleotide diversity among conserved domains (5, 14, 16). These characteristics position the 42 kDa region as a prime vaccine candidate.

The allelic diversity of *pkmsp1* in Thailand is influenced by intragenic recombination events within the *pkmsp1_42_*, as indicated by the presence of potential recombination sites and a decrease in linkage disequilibrium (LD) with increasing nucleotide site distance. Analysis of the haplotype network of *pkmsp1_42_* illustrates the relationship between parasite populations from different time periods. It shows that most haplotypes in the recent population originated from previous populations, with few new haplotypes emerging. This suggests that recent *P. knowlesi* strains circulating in Thailand are descendants of those from previous decades, with surviving variants exhibiting higher fitness. The haplotype network also did not show distinct clustering between different sampling periods, indicating a continuous relationship between parasites from recent and former periods. Thus, parasites from the recent population appear to be indigenous, originating from their ancestral parasites in the former population. Four haplotypes shared between Thailand and Malaysia signify the historical common origin of specific variants across these locations. This genetic exchange could occur through the movement of infected humans, monkeys, or vectors between regions. Notably, haplotype 3 (H3) exhibits the widest transmission range, spanning from Chantaburi in Thailand to Kelantan, Pahang, and Sabah in Malaysia. This extensive distribution suggests a dispersal of this particular genetic variant across different geographic areas. Similarly, haplotype 6 (H6) shows a broad transmission pattern, being identified in Narathiwat of Thailand and Terengganu, Selangor, and Sabah of Malaysia. In contrast, haplotypes 7 (H7) and 12 (H12) display a narrower transmission range, primarily crossing between southern Thailand and Peninsular Malaysia. This more localized distribution may indicate restricted movement of these variants, possibly reflecting localized transmission dynamics or geographical barriers influencing parasite spread.

The F_st_ value serves as a measure of genetic differentiation between populations. Generally, F_st_ values between 0.05 and 0.15 indicate low differentiation, while values between 0.15 and 0.25 signify moderate differentiation. Values above 0.25 indicate high differentiation (45). In our study, we observed significant genetic differentiation among parasite populations from different geographical regions. Specifically, there was a high level of genetic differentiation between parasite populations from central Thailand and southern Thailand (F_st_ = 0.3553) , as well as between central Thailand and Peninsular Malaysia (F_st_ = 0.3231). This finding is consistent with the expectation that geographical distance contributes to genetic divergence among parasite populations. Additionally, moderate genetic differentiation was observed between parasite populations from southern Thailand and Peninsular Malaysia (F_st_ = 0.1804), likely reflecting the shared landmass between these regions. It’s worth noting that our study did not distinguish between Malaysian isolates from Peninsular and Borneo due to potential sample size bias. However, previous research (45) has reported high genetic differentiation between parasite populations originating from Peninsular Malaysia and Malaysian Borneo. Furthermore, we detected moderate genetic differentiation between parasite populations from Thailand and Peninsular Malaysia. These findings together support the role of geographical factors in shaping the genetic diversity and differentiation of *P. knowlesi* populations in Southeast Asia.

However, this study has two principal limitations. Firstly, the analysis of multiclonal infections was constrained by the sequencing technology used, which only allowed for the inclusion of the major clone in each isolate. This approach may have obscured the underlying complexity of the infections, potentially leading to an underestimation of genetic diversity. Secondly, to ensure reliable analysis, only only high-quality sequences were retained, necessitating extensive trimming of the sequencing data. As a result, the obtained data only encompassed the 33 kDa portion of the 42 kDa region, with the 19 kDa segment excluded from analysis.

## CONCLUSIONS

A decline in genetic diversity based on *pkmsp1_42_* was evident amid the increased *P. knowlesi* incidence in Thailand over the past decade, suggesting rapid clonal expansion of the parasites. High synonymous substitution rates indicated purifying selection and a reduction in genetic diversity over time. Additionally, mutation points found on B-cell epitopes suggest the signature of host immune selection. Population differentiation analysis revealed a close relationship between the parasites in southern Thailand and Malaysia, indicating significant gene flow and recent shared ancestry in these connected border areas. These findings emphasize the need for coordinated efforts between Thailand and Malaysia to effectively address and eliminate *P. knowlesi* malaria.

## ACKNOWLEDGEMENTS

This research project is supported by the National Institute for Allergy and Infectious Diseases, The National Institute of Health (U19 AI089672 and U19 AI181583) and Mahidol University (Fundamental Fund: fiscal year 2024 by National Science Research and Innovation Fund (NSRF))

